# Aggregated Low-Density Lipoprotein (agLDL) Triggers Inflammatory Responses in Macrophages through Toll-Like Receptor 4 (TLR4)

**DOI:** 10.64898/2026.09.24.754244

**Authors:** Samia Kadri, Zeina Salloum, Judy Zhen, Cheng-I J. Ma, Dajiang Zhang, Thomas Lagace, Frederick R. Maxfield, Xiaohui Zha

## Abstract

Atherosclerosis is a chronic inflammatory disease primarily initiated by the sub-endothelial retention and modification of low-density lipoproteins (LDL), such as oxidation and aggregation. While oxidized LDL (oxLDL) exists in atherosclerotic lesions, aggregated lipoproteins are also prominent. Aggregated LDL (agLDL), for instance, is viewed as a passive substrate for macrophages to uptake and form foam cells. Recent studies show that agLDL, through toll-like receptor 4 (TLR4), induces formation of lysosomal synapses, which drive agLDL degradation by macrophages, a process described as “digestive exophagy”. As TLR4 is one of the pattern-recognition receptors for inflammatory responses in macrophages, we postulated that agLDL could initiate macrophage inflammation along with lipoprotein degradation/internalization, thereby contributing to a low degree of chronic inflammation in atherosclerosis. Here, we demonstrate that agLDL directly activates the canonical NF-κB signaling pathway and triggers a pro-inflammatory transcriptional program through TLR4, which closely resembles classical lipopolysaccharide (LPS)-mediated TLR4 activation. In addition, agLDL also activates TLR4 to generate inflammatory macrophages that recruit immune cells, a key process in the development of atherosclerotic lesions. Our findings therefore support a novel dual functionality of agLDL: a lipid donor and an inflammatory agonist. This establishes the agLDL-TLR4-NF-κB axis as a direct link between LDL aggregation in the intima and the initiation of sterile vascular inflammation. It also strengthens the understanding of the role of digestive exophagy with new mechanistic details.

---

Atherosclerosis is a chronic inflammatory condition of the arterial wall that represents a leading cause of global mortality^1^. The disease is initiated by high plasma LDL levels and subendothelial LDL accumulation. Trapped LDL then undergoes modification, such as oxidation and aggregation to form oxidized LDL (oxLDL) and aggregated LDL (agLDL), respectively. Macrophages subsequently digest these modified LDLs extracellularly and intracellularly to form lipid-laden foam cells, a process central to the progression of early atherosclerotic lesions. While the role of oxLDL in promoting oxidative stress and inflammation is known^2^, whether and how agLDL contributes to inflammation in atherosclerosis is much less clear. Aggregated lipoproteins, including agLDL, are prominent features in atherosclerotic lesions^3-5^. As such, agLDL may actively contribute to the non-resolving inflammation within the intima, in addition to its role as a lipid donor for foam cell formation.

A key development in recent years has been the identification of Toll-like receptor 4 (TLR4) as an essential mediator for agLDL degradation by macrophages. A series of seminal studies has established that agLDL requires macrophage surface TLR4 and MyD88 to form lysosomal synapses, resulting in agLDL digestion/endocytosis, release of fatty acids and cholesterol from agLDL, and subsequent foam cells formation^6-8^. Interestingly, TLR4 is a prominent pattern-recognition receptor that, upon engaging with ligands, mediates classical inflammatory responses in macrophages and, in the context of atherosclerosis, promotes lesion formation^9^. To date, little is known about the potential signaling consequences of TLR4 engagement with agLDL. Ligand-TLR4 interaction is well established to activate NF-κB and initiate inflammatory responses in macropahges^10^. However, agLDL has no known epitopes that could serve as canonical ligands to engage TLR4. Consequently, it remains unclear whether agLDL initiates the canonical NF-κB pathway to trigger inflammatory responses during its internalization. Resolving this question is vital to advancing our understanding of atherosclerosis and its accompanying lesion inflammation^1^.

In this study, we demonstrate, for the first time, that agLDL triggers potent inflammatory responses in macrophages through the canonical NF-κB signaling pathway. We provide evidence that agLDL induces a pro-inflammatory activation program that resembles classical lipopolysaccharide (LPS) stimulation. This inflammatory activation by agLDL is strictly dependent on TLR4 and is specific to the aggregated form of LDL. Importantly, agLDL exposure yields pro-inflammatory macrophages that can recruit immune cells, thereby functionally promoting atherosclerotic lesion development. These findings thus uncover a previously uncharacterized, major signaling axis in vascular inflammation, highlighting a critical pathway through which agLDL exacerbates atherosclerosis.

## Results

To determine whether agLDL could activate inflammatory responses in macrophages, we exposed bone marrow-derived macrophages (BMDMs) to agLDL (2.5 mg/ml) for 3 hours. AgLDL significantly increased the expression of several pro-inflammatory cytokines, including TNF-α, IL-1β, IL-6, and IL-12 (Fig. 1A). To determine whether agLDL also promoted inflammatory cytokine secretion, cytokine secretion from BMDMs exposed to agLDL was assessed. AgLDL induced robust secretion of TNF-α and IL-1β, which correlated with their gene expression (Fig. 1B). To ensure that the inflammatory activation was due to LDL aggregation and not LDL itself or a storage artifact, we exposed BMDMs to either agLDL or unmodified LDL prepared from the same LDL aliquot. Only agLDL, and not an equivalent amount of unmodified LDL, activated cytokine expression (Fig. 1C). We thus concluded that the aggregation of LDL was necessary for the inflammatory activation.

**Fig 1.**
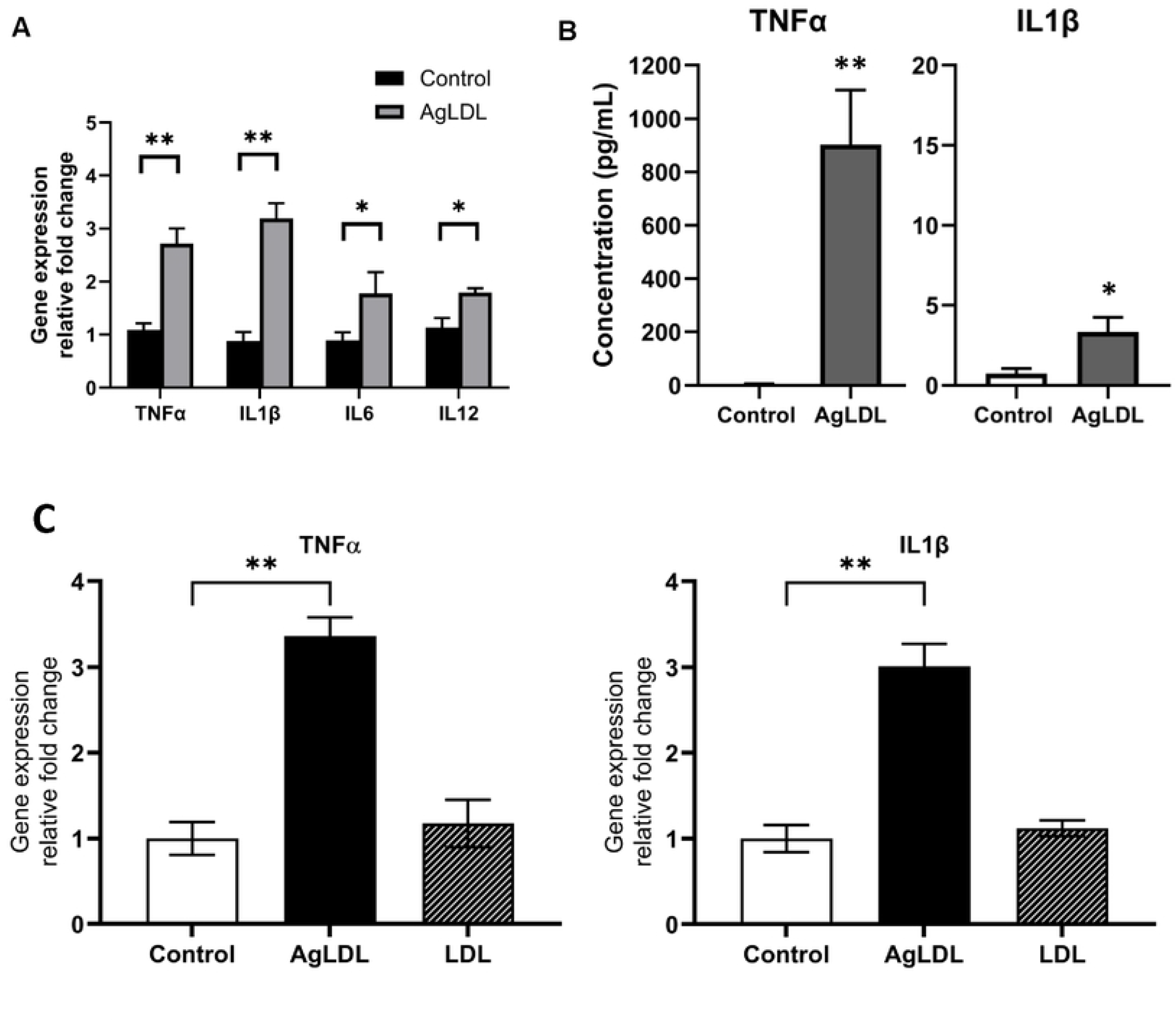
AgLDL triggers proinflammatory cytokine production in macrophages. **(A)** Bone marrow-derived macrophages (BMDMs) were exposed to or not exposed to agLDL (2.5 mg/ml) for 3 hours, and gene expression was analyzed by qPCR. **(B)** ELISA analysis of the expression of inflammatory factors, TNFα and IL-1β, after incubation with agLDL. **(C)** BMDMs treated with media only (control), agLDL, or LOL for 3 hours; TNFα and IL-1β mRNA expression was analyzed by qPCR. Data are representative of at least 3 independent experiments with 3 samples per group, and data are presented as mean ± standard deviation (SD). Statistical analysis was performed using an unpaired, two-tailed Student’s T-test. Asterisks (*), (**), and (***) indicate a significant difference with p < 0.05, p < 0.005, and p < 0.001, respectively.

We next examined the time course of agLDL-induced cytokine expression. We exposed BMDMs to agLDL (2.5 mg/ml) for 3, 6, and 9 hours, and quantified cytokine gene expression. Our results showed that TNF-α and IL-1β expression significantly increased linearly over time (Fig. 2A, B). We next characterized the dose-dependent effects of agLDL on inflammatory gene expression. We exposed BMDMs to increasing concentrations of agLDL for 3 hours and assessed pro-inflammatory cytokine expression. The expression levels of both TNF-α and IL-1β positively correlated with increasing agLDL concentrations (Fig. 2C, D). Overall, the inflammatory cytokine expression pattern is similar to LPS stimulated inflammatory responses.

**Fig 2.**
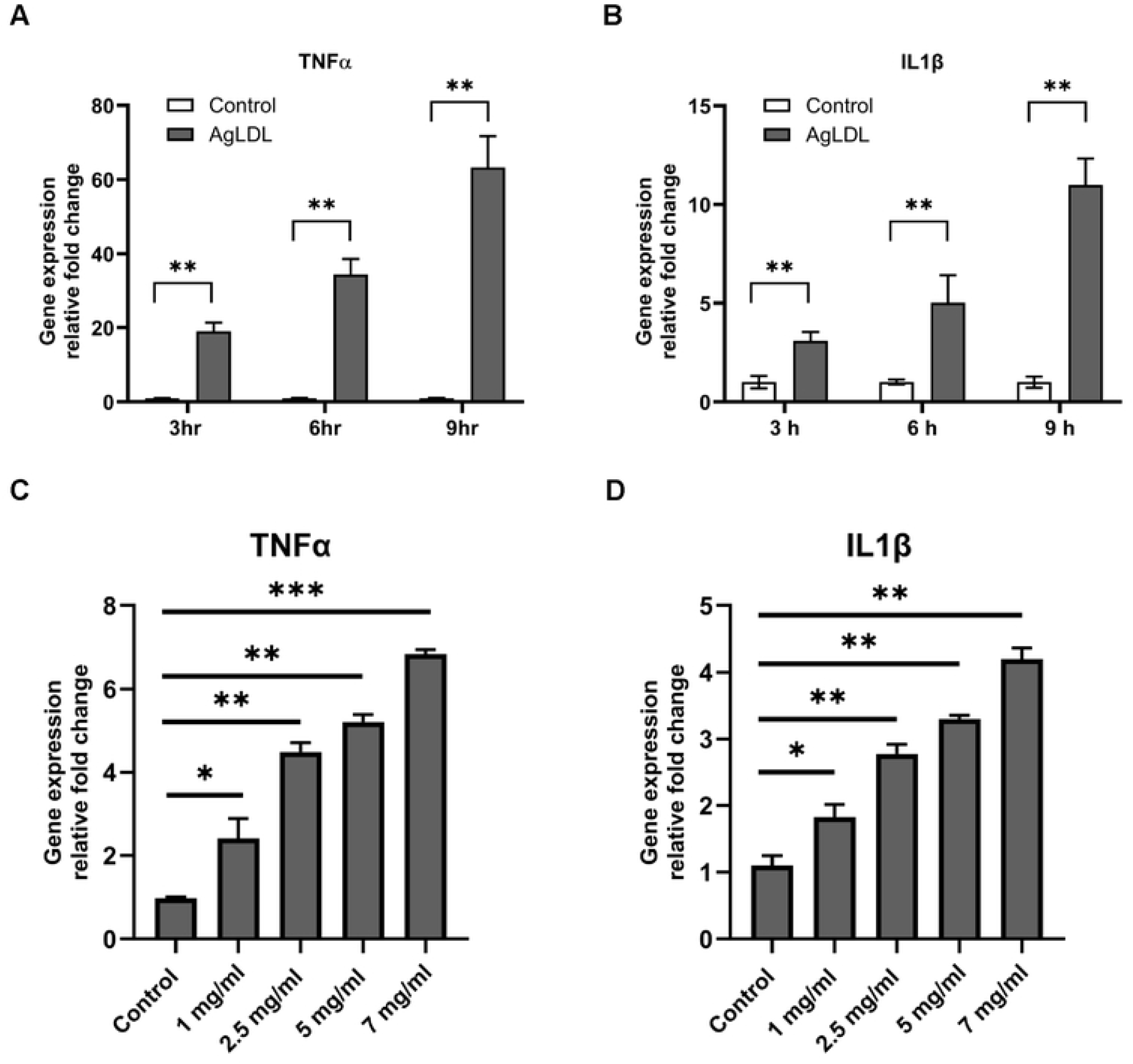
AgLDL stimulates a time-dependent and dose-dependent increase in proinflammatory cytokine expression and secretion. (A&B) BMDMs were exposed to or not exposed to agLDL at different time points, and TNFα (A) and IL1β (B) gene expression were analyzed. (C & D) BMDMs were exposed to different dosages of agLDL; TNFα (C) and IL1β (D) gene expression was analyzed by qPCR, and results were normalized to the control cells. Data are representative of at least 3 independent experiments with 3 samples per group, and data are presented as mean ± standard deviation (SD). Statistical analysis was performed using an unpaired, two-tailed Student’s T-test. Asterisks (*), (**), and (***) indicate a significant difference with p < 0.05, p < 0.005, and p < 0.001. respectively.

To understand how agLDL activated the inflammatory response in macrophages, we first tested whether agLDL activated the NF-κB pathway. As shown in Fig. 3A, agLDL increased the phosphorylation of the NF-κB p65 subunit (phospho-p65), nearly identically to activation by LPS.

**Fig 3.**
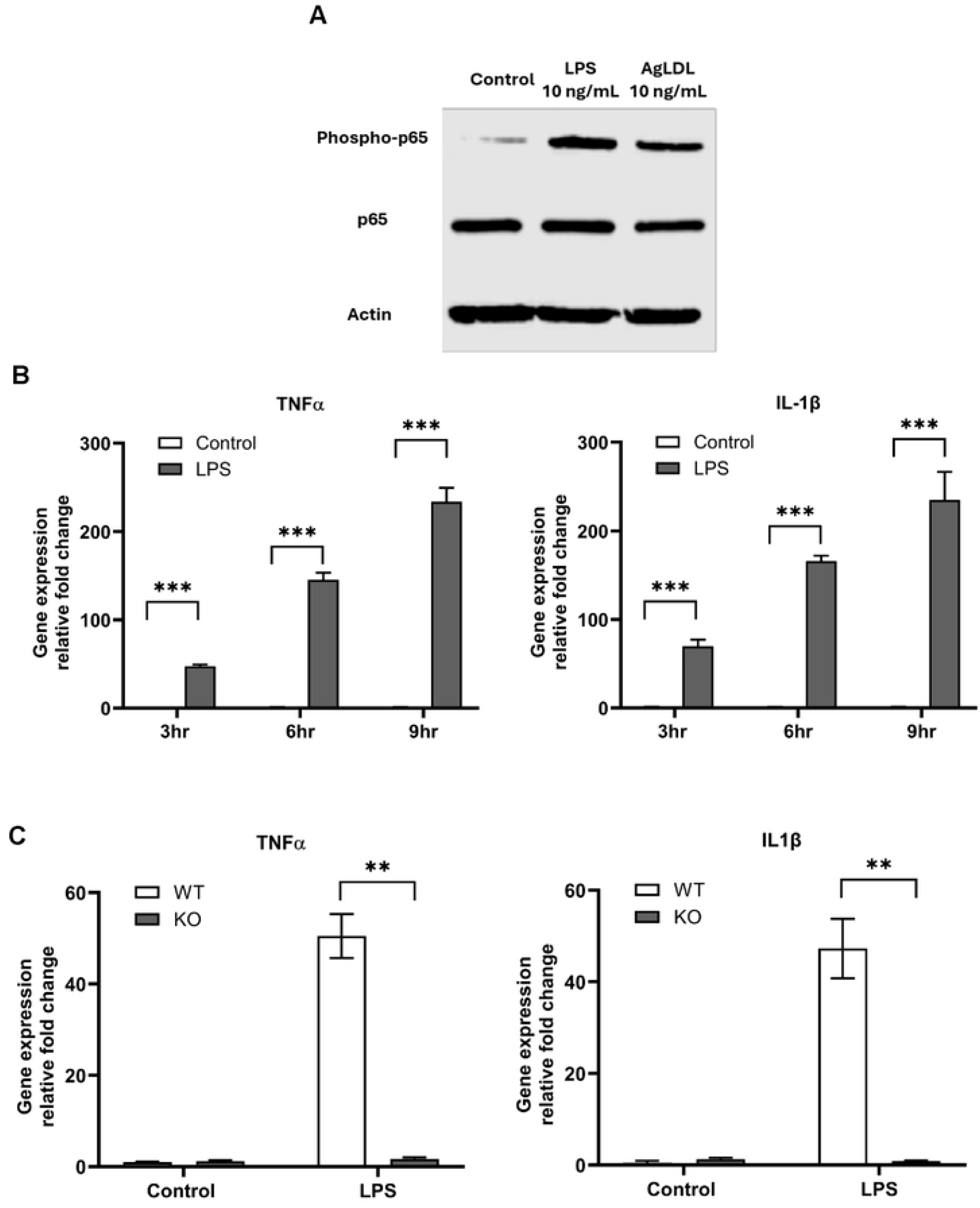
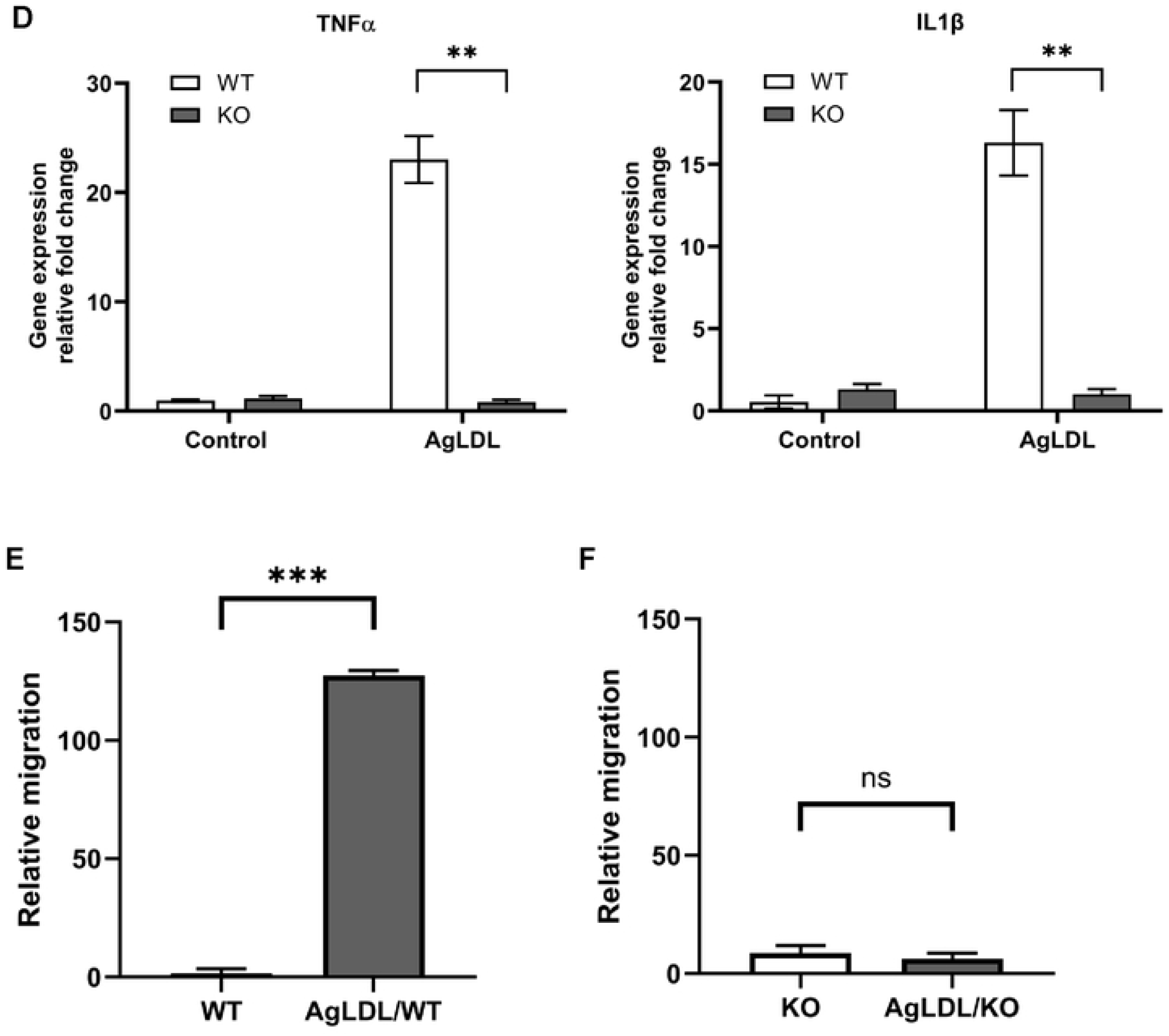
AgLDL induces activation of TLR4/NF-kB pathway. **(A)** Western blot of phosphorylated p65 subunit of the NF- **k**B (phospho-p65) protein complex in BMDMs stimulated with LPS or AgLDL for 3 hrs. Band intensities were quantified by densitometric analysis using ImageJ. Protein expression of phospho-p65 was measured relative to β-actin used as an internal control. (B) TNFα and IL1β gene expression in BMDMs, incubated with or without LPS (10 ng/ml) for different periods, was analyzed by qPCR. **(C&D)** Wild-type BMDMs incubated with or without LPS (C) or AgLDL (D) for 3 hr, and gene expression of TNFα and IL-1β was analyzed by qPCR. **(E & F)** Transwell migration assay ot TLR4-KO (E) and WT (F) BMDMs treated with or without agLDL for 3 hrs. Cell migration was quantified and is presented as the relative mmber of migrated cells. Data are representative of at least 3 independent experiments with 3 samples per group, and data are presented as mean ± standard deviation (SD). Statistical analysis was performed using an unpaired, two-tailed Student’s T-test comparing each treatment or condition to cells alone. Asterisks (*), (**), and (***) indicate a significant difference with p < 0.05, p < 0.005, and p < 0.001, respectively.

We then compared the temporal response of macrophages to LPS with the response to agLDL. LPS activated the expression of TNF-α and IL-1β linearly over 9 h (Fig. 3B), matching the pattern observed with agLDL (Fig. 2A, B). These data strongly suggested that agLDL might activate the same signaling pathway as LPS. LPS exclusively binds to TLR4 to initiate inflammatory responses^11^. Interestingly, the digestion of agLDL by macrophages requires TLR4 on the cell surface^7^. It is therefore possible that agLDL might require TLR4 to activate the inflammatory response. We therefore tested the effects of agLDL in TLR4-deficient macrophages. We isolated BMDMs from wild-type and TLR4-deficient mice, exposed them to either LPS or agLDL, and then analyzed inflammatory cytokine expression. As expected, the expression of TNF-α and IL-1β in response to LPS was strictly TLR4-dependent (Fig. 3C). Similarly, agLDL only activated TNF-α and IL-1β expression in wild-type BMDMs, whereas no activation was observed in TLR4^-/-^ BMDMs (Fig. 3D). We therefore concluded that agLDL stimulated inflammatory responses in macrophages by activating TLR4 and its downstream signaling pathway.

So far, we have demonstrated that agLDL induced macrophages to express and secrete inflammatory cytokines. However, compared to LPS, the inflammatory responses elicited by agLDL were much milder. Nevertheless, such activation levels by agLDL may be physiologically relevant in the context of atherosclerosis, where low-grade inflammation is predominant^1^. We thus examined whether agLDL could exert a significant pathological impact. In atherosclerosis, a primary function of macrophage inflammatory activation is to recruit immune cells^12,13^, which directly contributes to lesion formation. In this context, it was essential to determine whether the cytokine secretion from agLDL-exposed macrophages was biologically relevant and capable of recruiting immune cells, and whether this process was TLR4-dependent. We therefore performed transwell migration assays where BMDMs seeded in the bottom chamber were exposed to agLDL. We then analyzed BMDM downward migration from the top filter chamber. As demonstrated here, when BMDMs in the bottom chamber were exposed to agLDL, the downward migration of cells from the top chamber significantly increased, compared to cells alone (Fig. 3E). Furthermore, this migration was strictly TLR4-dependent; TLR4^-/-^ BMDMs exposed to agLDL completely failed to induce downward migration (Fig. 3F), which correlated with the absence of expression of inflammatory cytokines in these knockout cells (Fig. 3D).

We therefore conclude that agLDL induces TLR4-mediated inflammatory activation in macrophages, which is directly relevant to atherosclerosis pathology. AgLDL generates inflammatory macrophages to recruit additional immune cells, a key process in the development of atherosclerotic lesions. Thus, LDL aggregation could directly contribute to vascular inflammation and lesion formation during atherosclerosis, in addition to delivering lipids for foam cell formation.

## Discussion

Atherosclerosis is fundamentally an inflammatory lipid storage disease of the arterial wall^1^. In this study, we demonstrate for the first time that aggregated LDL (agLDL) directly triggers a robust pro-inflammatory response in macrophages through TLR4 and the canonical NF-κB signaling pathway. Although lipoprotein oxidation is a common modification that occurs in vivo and contributes to several atherogenic processes, it cannot account for all aspects of atherogenesis, particularly macrophage foam cell formation^14^. Aggregated lipoproteins are abundant in atherosclerotic lesions, arising from LDL particles that become trapped within the arterial intima and subsequently aggregate^4,5^. Macrophages respond to agLDL by forming extracellular hydrolytic compartments, or lysosomal synapses, in a TLR4-dependent manner, enabling extracellular digestion and lipid uptake that promotes foam cell formation^15^. Our findings here reveal an additional pathological consequence of agLDL. Beyond serving as a substrate for lipid accumulation, agLDL directly activates macrophage inflammatory signaling through TLR4 and stimulates chemokine production that recruits immune cells. Thus, agLDL is not merely a passive source of lipid but an active driver of chronic arterial inflammation, providing a mechanistic link between lipid deposition and innate immune activation during atherogenesis.

The mechanism by which agLDL activates TLR4 remains unclear. Ultrastructural studies have shown that agLDL forms large, amorphous aggregates that enter deeply convoluted, surface-connected compartments of macrophages, now recognized as lysosomal synapses^6−8,15,16^. TLR4 is required for the formation of these specialized compartments in response to agLDL^7^, suggesting that lipid uptake and inflammatory signaling are mechanistically coupled. However, how agLDL and TLR4 cooperate to initiate both lysosomal synapse formation and inflammatory activation remains an important unanswered question.

Given our results presented here, we propose the following model. Elevated plasma LDL accumulates within the arterial intima, where it becomes trapped, aggregated, and is likely anchored to extracellular matrix components^1^4. As professional scavengers, macrophages would attempt to engulf these large aggregates. However, unlike conventional phagocytic targets, agLDL is too large, irregularly shaped, and likely immobilized within the extracellular matrix to undergo complete phagocytosis^16^. We speculate that this initiates frustrated phagocytic attempt, which remodels the cortical actin cytoskeleton and associated plasma membrane, generating locally rigid membrane domains that promote the clustering and oligomerization of membrane proteins such as TLR4. TLR4 requires dimerization to activate downstream signaling through MyD88 and the canonical NF-κB pathway, initiating inflammatory gene expression^17^. Notably, MyD88 is also required for agLDL-induced exophagy^7^, further supporting a mechanistic connection between TLR4 signaling and lysosomal synapse formation. We therefore propose that TLR4 signaling subsequently further promotes the dynamic membrane rearrangements, which extends membrane ruffles around the aggregate, facilitating lysosomal synapse formation. Together, these coordinated processes drive both extracellular digestion of agLDL and inflammatory activation.

Several observations support this model. Receptor-mediated phagocytosis of glass beads has seen to remodel the plasma membrane and actin cytoskeleton, thereby enhancing LPS-induced TLR4 signaling in macrophages^18^. Conversely, TLR4 activation promotes plasma membrane dynamics and increases macropinocytosis^19^, processes that could facilitate lysosomal synapse formation during agLDL induced formation of a lysosomal synapse. These findings suggest that agLDL degradation and TLR4 activation are two mutually reinforcing processes, with each promoting the other. Such reciprocal relationship may represent a previously unrecognized form of inflammatory digestive exophagy that integrates extracellular digestion with innate immune signaling.

More broadly, protein aggregate-induced TLR4 activation coupled to exophagy may represent a common mechanism in chronic inflammatory diseases beyond atherosclerosis. For example, amyloid plaques in Alzheimer’s disease may similarly trigger this process in microglia^20,21^. Consistent with this possibility, amyloid-β activates inflammatory signaling through TLR4- and MyD88-dependent pathways^22^. If confirmed, this mechanism would identify a general strategy by which large extracellular protein aggregates activate innate immune responses in diverse pathological settings.

Overall, our findings suggest that large extracellular protein or lipoprotein aggregates may represent a distinct class of endogenous danger signals. Rather than functioning through conventional ligand–receptor recognition, these aggregates may activate innate immune responses by remodeling the plasma membrane and promoting TLR4 clustering and activation. If this mechanism proves to be broadly applicable, it could provide a unifying framework for understanding inflammatory responses associated with diverse protein aggregation disorders, including atherosclerosis, Alzheimer’s disease, and other chronic degenerative diseases. More broadly, these findings highlight the importance of the physical properties of extracellular aggregates, in addition to their biochemical composition, in regulating innate immune signaling.

## Acknowledgements

This work was supported by a grant (grant-in-aid) from Heart and Stroke Foundation of Canada (HSFC), G-25-0041100, and a grant from Canadian Institute of Health Research (CIHR), PJT-180504.

## Experimental procedures

### Cells and cell culture

Bone marrow-derived macrophages (BMDMs) were differentiated from bone marrow cells isolated from the femora and tibiae of 4- to 16-week-old wild-type C57BL/6J or Tlr4 KO (B6(Cg)-*Tlr4*^*tm1*.*2Karp*^/J) mice. Mice were euthanized using CO_2_ gas asphyxiation, followed by cervical dislocation^23^. Then, long bone dissection and bone marrow isolation were performed as previously described^23^. BMDM preparation followed the protocol previously described^24^, with minor modifications^25^.

After filtration of the bone marrow isolate, the filtrate was centrifuged at 480 g for 10 minutes at room temperature. The supernatant was discarded, and the cell pellet was briefly (< 2 minutes) resuspended in red blood cell lysis buffer. The differentiation medium (DMEM + 20% L-929-conditioned media + 10% FBS + 1% penicillin-streptomycin) was added to the cell suspension, then the cells were centrifuged again at 480 g for 10 minutes at room temperature. The supernatant was discarded, and the cell pellet was resuspended in differentiation medium. The cell suspension was then filtered through a 70-um strainer into a 50mL centrifuge tube. The cells were then seeded into 100-mm diameter dishes (Greiner Bio-One, Monroe, NC) with a total volume of 10mL at 0.6-0.8 × 106 cells/mL (∼10,000-15,000 cells/cm^2^). The dishes were then incubated for 6 days at 37°C in a humidified atmosphere containing 5% CO_2_. At day 3 or 4, 10mL of differentiation medium was added to the dishes.

At the end of day 7, the cells were detached using a glass pipette, counted and reseeded at 0.5 × 106 cells/mL in 24-well plates. The medium used for reseeding at this step is DMEM supplemented with 10% FBS and 1% penicillin-streptomycin. Plates were incubated overnight at 37°C in a humidified atmosphere containing 5% CO_2_. On day 8, BMDMs were treated at the indicated time points with LDL, AgLDL, or LPS prepared in testing media composed of DMEM supplemented with 4 mM glutamine.

### Reagents and chemicals

Dulbecco’s Modified Eagle Medium (DMEM) was purchased from Gibco (Thermo Fisher Scientific, Watham, MA). FBS (Optima) was purchased from Atlanta Biologicals (R&D systems; Minneapolis, MN). Antibiotics (penicillin and streptomycin), fatty acid-free bovine serum albumin (BSA), and phosphate-buffered saline (PBS) were purchased from Sigma-Aldrich (St Louis, MO).

### Lipoproteins

LDL was supplied by the Lipoprotein Receptor Biology Laboratory at the University of Ottawa Heart Institute. LDL was isolated from fresh human plasma by sequential potassium bromide flotation ultracentrifugation followed by extensive dialysis against PBS containing 0.25mM EDTA^26^. The protocol for aggregating LDL was adapted from previous agLDL studies^7^. Aggregates of LDL were formed by vigorously vortexing LDL aliquots for 60 seconds. The agLDL was pelleted by centrifuging the tubes at 10,000 g for 10 minutes at 4°C. To isolate the agLDL, the supernatant was aspirated, and the pellet was resuspended in serum-free DMEM.

### RNA purification and cDNA synthesis

At the end of the incubation, the cells were lysed in TRIzol (ThermoFisher Scientific, Watham, MA). The cell lysate was collected and stored at -80°C until total RNA extraction. Phenol-chloroform extraction was performed as described in the TRIzol user guide. RNA was then purified using a Molecular Biology kit (BioBasic Inc., Markham, ON), according to the manufacturer’s instructions. RNA concentrations and RNA purity were measured using a NanoDrop One (ThermoFisher Scientific, Watham, MA) spectrophotometer. Prior to cDNA synthesis, RNA was stored at -80°C. cDNA synthesis was performed using a QuantiTect Reverse Transcription kit (Qiagen, Germantown, MD), following the manufacturer’s instructions. To this end, a Bio-Rad T100 PCR Gradient Thermal Cycler was used. cDNA was then stored at - 20°C.

### Quantification of gene expression

Gene expression was quantified using reverse transcriptase quantitative PCR (RT-qPCR). RT-qPCR was performed according to the Fast SYBR Green protocol using the Rotor-Gene Q PCR detection system (Qiagen, Germantown, MD). The genes analyzed were HPRT1, TNF-α, IL-1β, and IL-6. HPRT1 was used as a reference gene, while TNF-α, IL-1β, and IL-6 were the target genes. The primers were ordered from Invitrogen (ThermoFisher Scientific, Watham, MA), and the sequence for each primer is listed in Table 1^23^. For each sample and gene analyzed, technical triplicates were loaded.

**Table 1.**
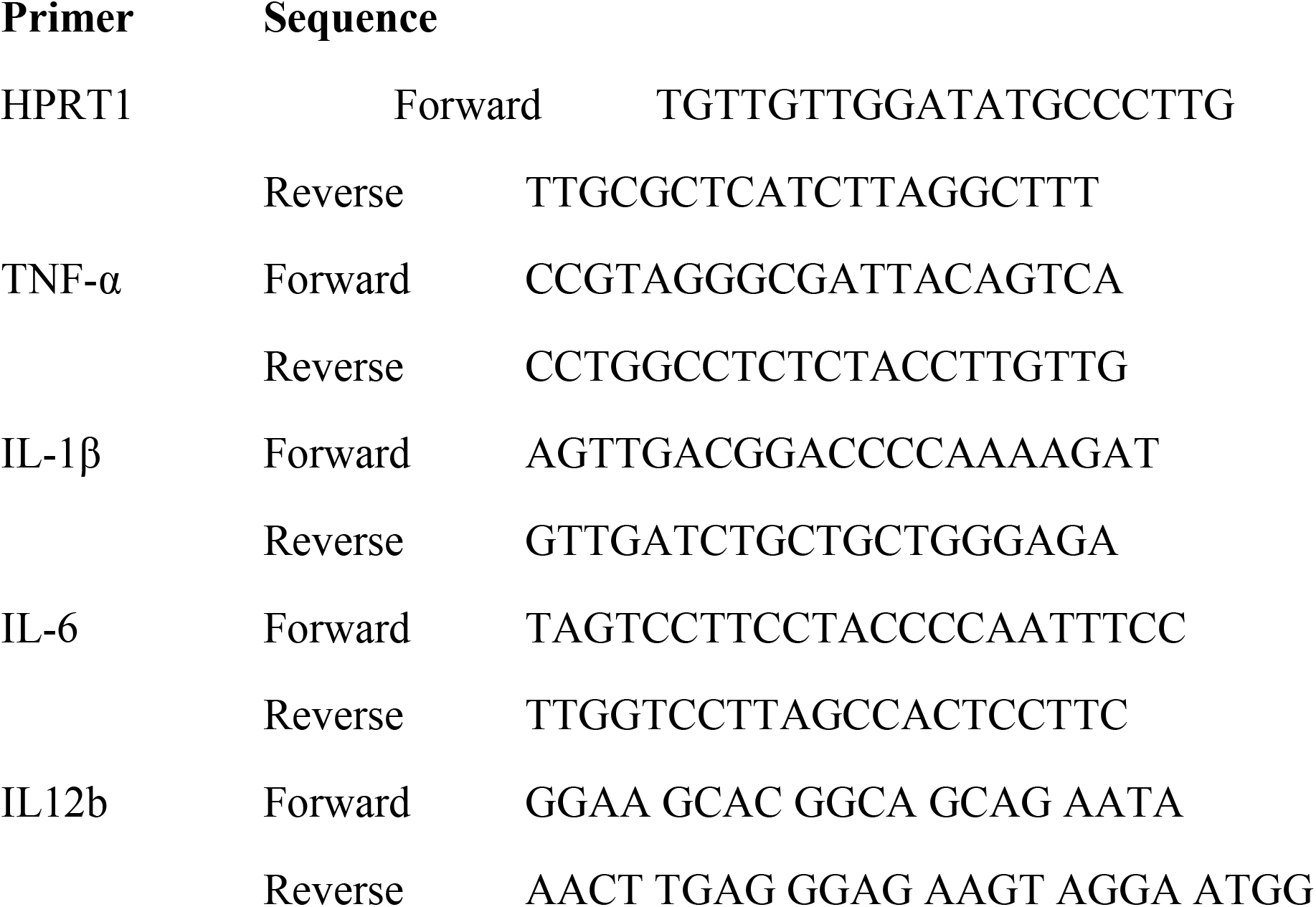
Mouse primers used for RT-qPCR.

### Cytokine ELISA Assay

At the end of the incubation, cell supernatants were collected and cell debris were removed by centrifuging at 12,000 × g for 5 min. Sample aliquots were stored at - 80°C until analysis. ELISA was performed using the DuoSet ELISA mouse TNF-α kit (R&D Systems; cat#DY410) to quantify TNF-α secretion, and the DuoSet ELISA mouse IL-1β kit (R&D Systems; cat#DY401-05) to quantify IL-1β secretion, according to the manufacturer’s instructions.

### Immunoblot analysis

At the end of the incubation, the cells were harvested and washed three times with PBS. After resuspension in lysis buffer (50 mM Tris-HCl, pH 7.5, 150 mM NaCl, 0.1% Triton-X 100, and protease inhibitor), the cells were kept on ice for 30 min, followed by centrifugation at 500 × g for 5 min. Total cell lysates (20 μg) were separated on 10% SDS-PAGE and subsequently electrotransferred onto nitrocellulose membranes. The membranes were incubated in 5% non-fat dried milk in Tris-Buffered Saline and Tween-20 (TBST; 50 mM Tris-HCl, pH 8.0, 150 mM NaCl, and 0.05% Tween-20) for 1 hr at room temperature. The blots were probed with one of the following primary antibodies diluted as indicated: NFkB p65 polyclonal antibody (1:1,000), phospho-NFkB p65 (Ser536) monoclonal antibody (1:100), and anti-actin mouse polyclonal antibody (1:1,000) at 4°C overnight. The membranes were washed with TBST and further probed with horseradish peroxidase-conjugated (HRP) anti-mouse or anti-rabbit IgG antisera (1:10,000) at room temperature for 1 hr. Following two washes with TBST, the specific proteins were visualized with a chemiluminescence HRP Substrate system (Millipore, Massachusetts, USA) using LAS 4000 (Fujifilm Life Science, Cambridge, USA) according to the manufacture’s protocol.

### Migration assay

Macrophage migration was assessed using 24-well Transwell inserts with 3-µm pore PET membranes (6.5-mm diameter; VWR, 10769-240). After 7 days of differentiation, BMDMs were resuspended in DMEM containing 10% FBS and 1% penicillin/streptomycin, and 10^5 cells were seeded onto each insert and incubated overnight at 37 °C and 5% CO_2_. Where indicated, 10^6 BMDMs were seeded in the lower chamber. The following day, cells were washed with prewarmed DMEM, and inserts were transferred to wells containing 400 µl DMEM with or without AgLDL (2.5 mg/ml) and with or without BMDMs in the lower chamber. Cells were allowed to migrate for 7–8 h at 37 °C and 5% CO_2_.

Following migration, inserts were washed with PBS and fixed in 4% paraformaldehyde for 10 min, followed by 70% ethanol for 10 min. Membranes were stained with 0.2% (w/v) crystal violet in 10% ethanol for 10 min, and the remaining cells on the membrane upper surface were removed with a cotton swab. Migrated cells (membrane lower surface) were quantified by solubilizing membrane-associated crystal violet in methanol and measuring absorbance at 570 nm. The assay was adapted from Justus et al.^27^.

### Statistical Analysis

Statistical analysis was performed using GraphPad PRISM software. The statistical significance of the differences between groups was analyzed using an unpaired Student’s T-test with Welch’s correction. Differences were considered significant when the p-value < 0.05.

## Notes

### Competing Interest Statement

The authors have declared no competing interest.

